# A frog adaptive radiation: Ecomorphological evolution in Old-World shrub frogs (Rhacophoridae: *Pseudophilautus*)

**DOI:** 10.1101/2023.08.12.549244

**Authors:** Madhava Meegaskumbura, Gajaba Ellepola, Gayani Senevirathne, Kelum Manamendra-Arachchi, Nayana Wijayathillaka, Marcio Pie, Dan Sun, Rohan Pethiyagoda, Christopher J. Schneider

## Abstract

Ecomorphs result from divergent natural selection, leading to species-rich adaptive radiations. Identifying ecomorphs and the resulting adaptive radiations in frogs is challenging due to conserved morphology and high species diversity. In this study, we demonstrate the ecological and climate specializations that have driven the diversification of shrub frogs of the genus *Pseudophilautus* in Sri Lanka, a tropical continental island. We use a time-calibrated phylogeny, morphometric analyses, and climate-niche evolution, and identify five ecomorphological categories, including Tree-shrub, Rock-boulder, Leaf-litter, Habitat Generalists, and Canopy forms, and describe their evolution. Body size is the primary factor separating species, and specific body features correlate with habitat type. Ecomorphs likely evolved multiple times in disparate lineages, and in different regions and altitudes, during cold climatic periods owing to monsoon cycles resulting from the Himalayan-Tibetan orogeny. The common ancestor was a medium-sized, wet-adapted, tree-shrub habitat specialist which originated in the late Oligocene. Extreme size classes (diminutive leaf litter forms and large canopy forms) evolved recently and suggest that morphological disparity arose late in diversification, possibly aided by favorable climates. This work will facilitate understanding of adaptive radiations in frogs, which possibly will help uncover the prevalence of subtle adaptive radiations in frogs, just as in tailed-vertebrates.

## Introduction

Adaptive radiations play a pivotal role in the phenotypic and ecological diversification of life on Earth. This occurs when species swiftly evolve from a single ancestral lineage into a myriad of distinct forms. Such rapid diversification often transpires when environmental changes present new resources, modify biotic interactions, or unveil previously unoccupied ecological niches [1](Schluter, 2000). The foundations of this process were established by [2]Darwin, (1845) and refined by generations of evolutionary biologists, providing valuable insights into the evolutionary patterns and causal processes seen in various groups of organisms [3-12](Glor, 2010; Losos & Miles, 2002; Ole, 2006, Losos & Ricklefs, 2009; Losos & Mahler, 2010, Blackburn et al. 2013, Close et al. 2015, Tokita et al. 2017, Tiago et al. 2020, Swardal et al. 2020). The ecological theory of adaptive radiation suggests that speciation in response to divergent natural selection is primarily ecological [1,3,6](Glor, 2010; Losos & Mahler, 2010; Schluter, 2000). However, studying the processes that lead to ecological speciation is challenging for long-lifespan species, such as vertebrates. Hence, adaptive radiations are typically studied as a macroevolutionary process in deep time, by evaluating the signatures of past or ongoing ecological speciation processes that resulted in reproductive isolation [3,13-15](Glor 2010; Astudillo-Clavijo 2015; Arnold et al. 2008; Revell et al. 2008).

Ecomorphological clustering is a hallmark of adaptive radiations spurred by ecological speciation. It denotes the interconnected relationship between the physical characteristics (morphology) and ecological roles of closely related species [16,17](Aerts et al. 2000, Schluter, 2001). Ecomorphological clustering is an essential indicator of past or ongoing ecological speciation processes and includes the existence of ecomorphs and divergence in morphology and ecology among sister species [18,19](Erwin, 1994, Moen et al. 2021). In addition to ecomorph evolution, adaptive radiations typically exhibit an ‘early burst’ pattern, where speciation rates surge in the initial stages of radiation and then decline as ecological niches become saturated over time [4,18,20](Simpson, 1953; Erwin, 1994; Losos and Miles, 2002).

Frogs and toads (Anura) are one of the most species-rich tetrapod orders with about 8000 species [21](AmphbiaWeb, 2023), yet their adaptive radiations are not well understood. Anurans have a conserved body plan [22](Duellman & Trueb, 1986) which results in fewer opportunities for adaptations compared to tailed vertebrates that often display remarkable adaptive radiations. However, frogs can still undergo adaptive radiation and have ecomorphs, although it is more subtle compared to other species [7,23] (Bossuyt and Milinkovitch 2000, Blackburn et al. 2013).

The species-rich shrub-frog of the genus *Pseudophilautus* [24](Meegaskumbura et al. 2002) characterized by direct development, a key evolutionary innovation that allows terrestrial niche occupation [25](Meegaskumbura et al. 2015), provides an opportunity to investigate whether diversification in these frogs constitutes of ecomorphs indicative of ecological speciation.

The ancestor of *Pseudophilautus* dispersed to Sri Lanka across the ephemeral Palk Isthmus, which provided a terrestrial connection with India during periods of lowered sea level [26](Sudasinghe et al. 2018). Following initial diversification in Sri Lanka, a small component of these frogs back-migrated to India [27,28](Bossuyt et al. 2004, Meegaskumbura et al. 2019). The species remained occupy a diversity of habitats [29-35](Manamendra-Arachchi and Pethiyagoda 2005; Meegaskumbura and Manamendra-Arachchi 2005; Meegaskumbura et al. 2007, 2009, 2012; Wickremasinghe et al. 2013; Meegaskumbura and Manamendra-Arachchi 2011), mostly in the island’s aseasonal southwestern wet zone. The wet zone includes a large area of lowland rainforest and three isolated mountain massifs, which combine to provide a climatically and topographically complex region with substantial habitat heterogeneity. Within the wet zone, the vast majority of species are confined to the few remaining fragments of rainforest and tropical montane cloud forest, where they occupy a variety of microhabitats, including rock crevices and boulders in streams, leaf litter, shrubs, open grasslands, and the low, mid- and high canopy of trees [29,30,32,33](Manamendra-Arachchi and Pethiyagoda 2005; Meegaskumbura and Manamendra-Arachchi 2005, 2007, 2009, 2012).

We hypothesize that the ecologically disparate habitat occupation and specialization of *Pseudophilautus* species signals the existence of ecomorphs that underlie a spectacular adaptive radiation. To document such ecomorphs (phenotype-habitat correlation) and outline their evolution, we analyze this species-rich, monophyletic and predominantly insular radiation of tree frogs using a multi-gene, time-calibrated phylogeny, ancestral-state reconstructions, and morphometric analyses. We predict that body size and shape may be influential in explaining the phenotype-habitat correlation of *Pseudophilautus*, and that historical climate may have played a critical role in its diversification.

## Results

### Ecological specialization

Our study of Sri Lankan *Pseudophilautus* showed that these frogs can be classified into five distinct ecomorph categories: Canopy, Generalist, Rock-boulder, Leaf-litter and Tree-shrub in the morphospace (Fig.1, Extended Data Figure 1). Using a principal component analysis (PCA; Fig. 1a), a phylogenetically corrected principal component analysis (pPCA; Extended Data Figure 1) and a linear discriminant analysis (LDA; Fig. 1b, 1c), we found that body size and shape can be used to differentiate between these categories (results of LDA analyses are provided in detail in Extended results). In the PCA, the first two principal components explain over 94% of the variance (Table S1, Extended Data Figure 2) with males and females showing a similar pattern within the morphological space. The first PC representing body size explains 93.2% of the morphological variance across species and appears to be the most important feature for separating the five clusters. Leaf-litter species have the smallest body sizes, while Canopy species have the largest. On the second PC axis, although the variance is low 1.5%), upper eyelid width (UEW) and palm length (PAL) load strongly mainly distinguishing Rock-boulder species from the others (Fig. 1A). The pPCA suggests that similar morphs have arisen from different lineages pointing towards a convergent evolution in *Pseudophilautus* (Extended Data Figure 1, Table S2).

**Fig. 1.**
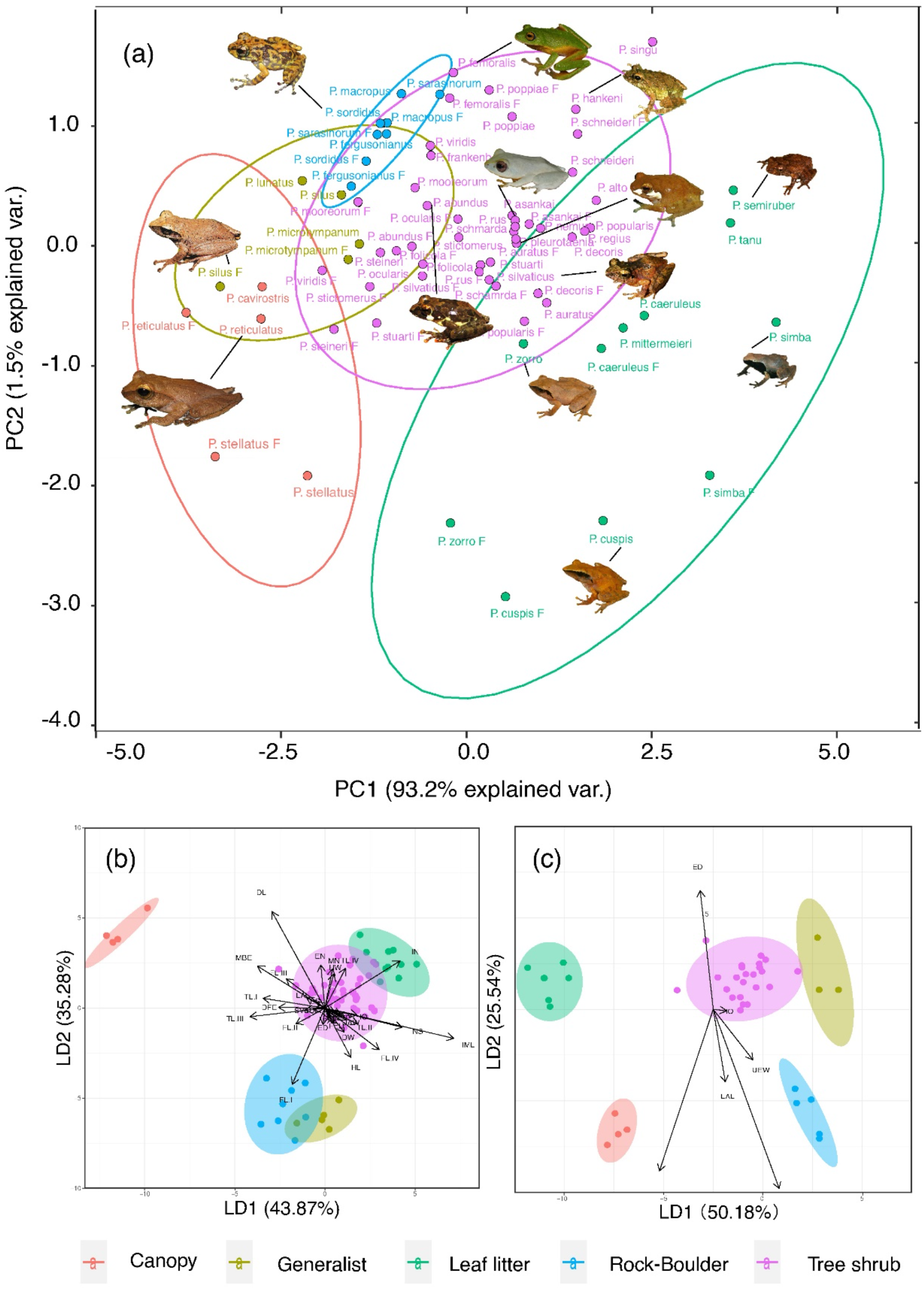
The position of Sri Lankan *Pseudophilautus* in morphological space as indicated by principal component (PCA—raw data) and linear discriminant analyses (LDA). The PCA conducted on raw data (a) separates species into five habitat categories along the PC1 axis, though with visible overlapping areas. Circles indicate the 95% confidence interval band for each habitat cluster. An ‘F’ following species names denotes females. Canopy forms and Tree-shrub forms are well separated in morphological space. The LDA conducted on raw data (b) groups all measurements into five habitat categories, with slight overlaps between Generalists and Rock-boulder forms and between Leaf-litter and Tree-shrub forms. Canopy forms are well separated from all others. The LDA conducted on size-corrected measurements (c) also groups species into five categories, with no overlap among groups. Head, finger and toe dimensions are decisive characters in defining their habitat correlation. Coloured circles indicate the 95% confidence interval band for each habitat cluster and each point represents a species. See Extended Data Figure 1 for the results of the pPCA analysis. Frog images indicate the relative size of each species.

The shape analysis based on size-adjusted measurements (residual data) in the PCA retains four principal components, which together explain 64.1% of the variation in shape (Table S3). Finger lengths (FL I-IV), foot length (FOL) and toe lengths (TL I-IV) load strongly on the first principal component. Head length (HL) and mandible–nostril distance (MN), inter-narial distance (IN), and distance between front of eyes (DFE) and inter-orbital width (IO) load strongly along PC2, PC3 and PC4, respectively. Although a distinct differentiation of shape isn’t observed among ecomorphs, Canopy forms are clearly separated from Generalists by having greater head length (HL) and mandible– nostril distance (MN) (Extended Data Figure 3, PC1 vs. PC2). Generalists can be distinguished from Rock-boulder forms and Tree-shrub forms by having shorter internarial distance (IN) (Extended Data Figure 3, PC1 vs. PC3). Canopy forms are separated from Rock-boulder forms by having longer fingers (FL), feet (FOL) and toes (TL) (Extended Data Figure 3, PC1 vs. PC4). However, Rock-boulder forms, Tree-shrub forms, Generalists and Canopy forms seem to exist within the ecomorphological space of Leaf-litter forms but, Tree-shrub forms deviate from this pattern along PC4 axis.

### Phylogenetic analyses

The phylogenetic analysis shows high node support for many taxa and the topology is consistent with [28] Meegaskumbura et al. (2019) (Fig. 2). The results suggest that the most recent common ancestor (MRCA) of the *Pseudophilautus* radiation evolved between 41.8 and 25.2 MYA, with major clade diversification occurring during early to mid-Miocene, 24.1–14.7 MYA. However, most of the extant species evolved during the late Miocene and the Pliocene. Lineage-through-time plots suggest an early, rapid speciation event γ = −3.7077 (p=0.04) over a brief period, which gradually slows down as new lineages accumulate with time (Fig. 2).

**Fig. 2.**
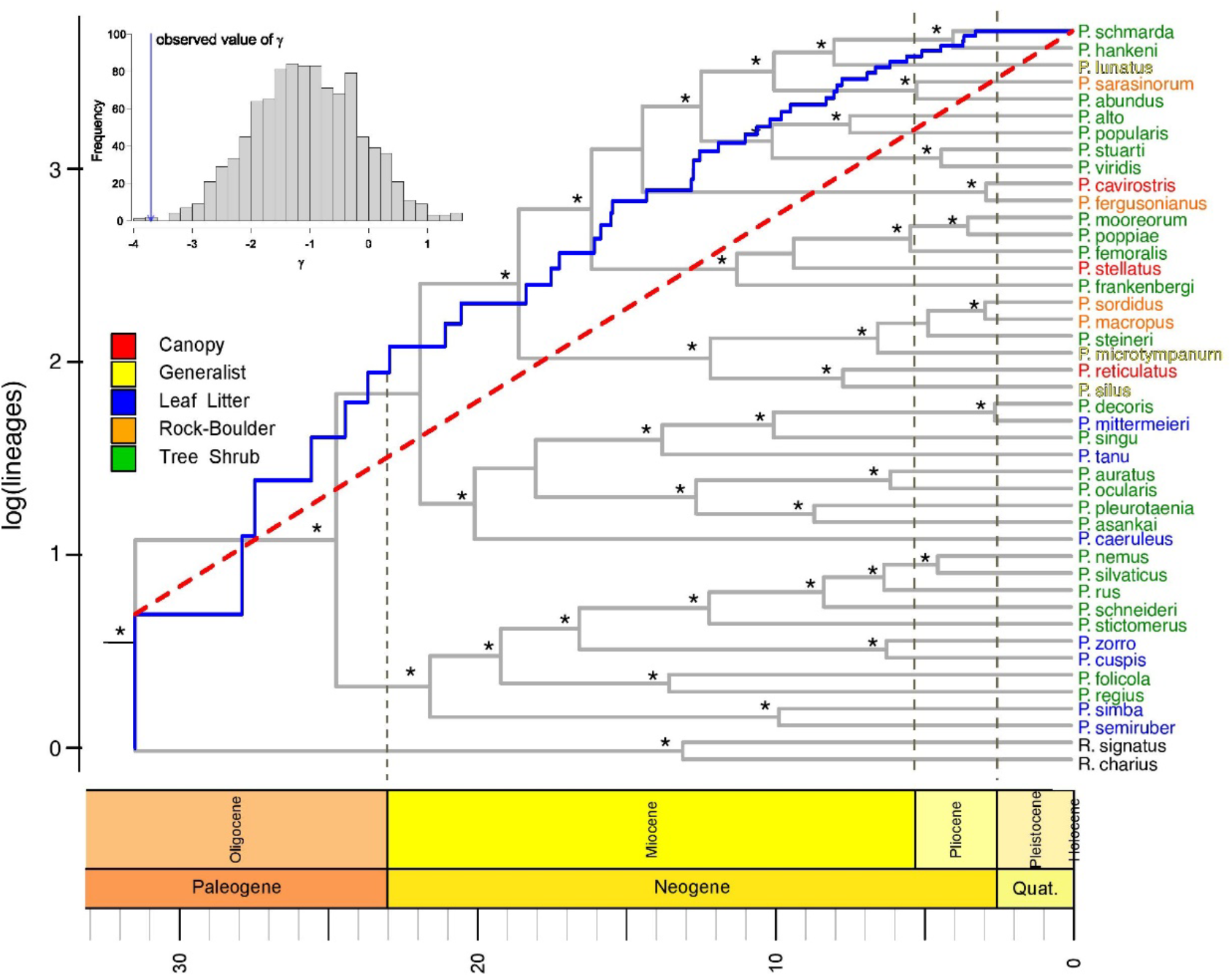
Rates of diversification and molecular phylogenetic relationships among extant *Pseudophilautus*, based on Bayesian inference of the concatenated data set of the 16S rRNA + 12S rRNA + CytB +Rag1 (2702 bp) loci. Asterisks (*) above nodes represent ≥ 95% Bayesian posterior probabilities. The blue line indicates the log lineage through time plot, whereas the null (Yule) model is indicated by the red dashed line. The lineage-through-time plot indicates an early burst of lineages (γ= - 3.707) followed by a decrease in speciation rate with time. The graph inset shows that having accounted for missing taxa, observed value of γ becomes significantly negative to the null model. Although speciation rates appear higher during the Miocene, Extant ecomorphs seem to have emerged during late Miocene and early Pliocene.

The MRCA of *Pseudophilautus* is predicted to be a medium-sized species (Fig. 3a). Early differentiation in body size (represented by PC1) accourred around 18.9 MYA, followed by an accelerated rate of body size evolution between 10 - 5 MYA, corresponding with the origin of ecomorph categories. Smaller-bodied frogs evolved to become Leaf-litter forms, while larger-bodied forms evolved to become Canopy dwellers, with a few reversals (Fig. 3a). However, there is no comparable trend in PC2 towards the evolution of ecomorphs (Extended Data Figure 3), indicating that body size evolution and evolution of morphological characters such as Upper eyelid width (UEW) and Palm length (PAL) were largely decoupled during the diversification of *Pseudophilautus* (Extended Data Figure 3).

**Fig. 3.**
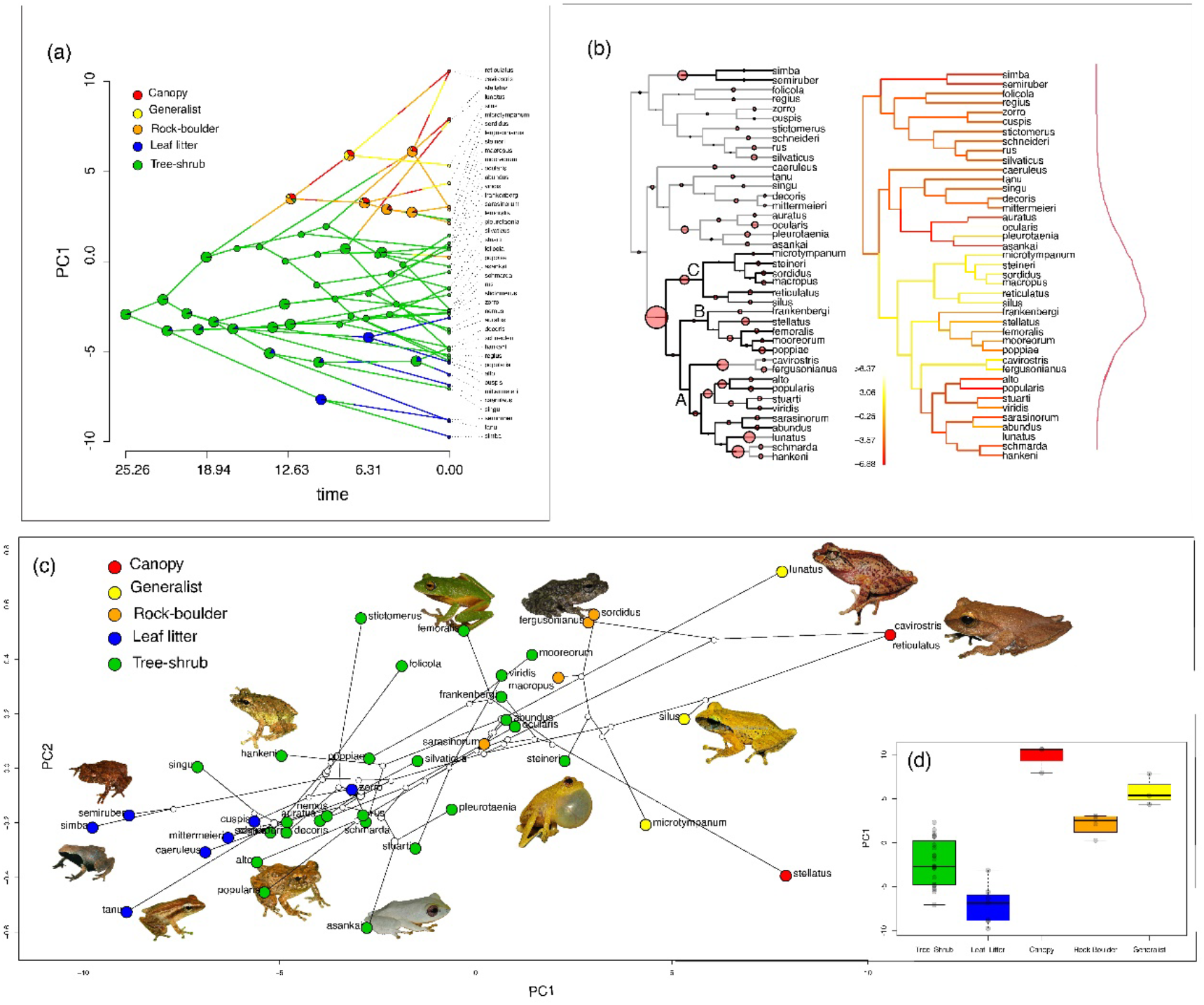
Temporal patterns of morphological evolution along PC1, evolutionary shifts in body size and relative position of ecomorphs in the phylomorphospace of *Pseudophilautus*. (A) Traitgrams of PC1 suggest that the different morphs evolved mainly by altering body size. Ancestral state reconstructions traced on traitgram (a) suggest that a medium-sized Tree-shrub form gave rise to the other ecomorphs, which likely evolved 10–5 MYA. (b) Most probable macroevolutionary rate shift configurations for the evolution of body size across the phylogeny. Circles on the left diagram denote shifts in rates of body size evolution. The most significant rate shift occurred near the crown of clades A, B and C. Generally, an initial shift towards larger body size was followed by shifts towards smaller body sizes, ultimately leading to a single adaptive optimum. Branch colors on the right diagram denote instantaneous rates (red = slow, yellow = fast). The probable number of adaptive peaks is denoted by the red curve alongside panel. (c) Different ecomorphs traced on the phylomorphospace indicates clear separation among the five ecomorphs along PC1, while Rock-boulder forms and Leaf-litter forms are clearly separated along PC2. (d) Box plots show separation of different ecomorph categories based on PC1 values (Table S4).

Interestingly, the ‘bayou’ analysis detected evolutionary rate shifts towards a single major adaptive peak (Fig. 3b; the red curve on right). An initial shift towards large body sizes (Fig 3b; large circle on the phylogeny denote rate shifts towards larger body sizes) was followed by shifts towards smaller sizes. The most significant rate shift coincided with the origin of clades A, B and C around 20 MYA (Fig. 2). These clades gave rise to derived ecomorphs such as Canopy forms, Rock-boulder forms and Generalists; the other three clades contain only Leaf-litter forms and Tree-shrub forms.

The phylomorphospace (Fig 3c) show that species with small-to-medium body sizes belong to the Leaf-litter and Tree-shrub forms, whereas the large-bodied species belong to the Canopy forms. The phylogenetic MANOVA suggest that there is a significant difference (p = 0.0009) in the position of ecomorphs in the phylomorphospace, particularly along PC1 (Fig. 3d; Table S4). However, there is some overlap among adjacent ecomorphs. It’s worth noting that the evolution of body shape, as represented by size-corrected PC values, doesn’t correspond with ecomorph categories. The evolution of body shape was gradual throughout the history of *Pseudophilautus*, and Tree-shrub species span the entire range of body shapes (Extended Data Figure 4).

Based on the ancestral reconstruction of the five ecomorphs, it appears that the diversification of *Pseudophilautus* occurred through a Tree-shrub ancestor (Fig. 4a) This ancestor likely inhabited high montane regions (Fig. 4c). Rock-boulder forms arose three times from the common ancestor of [*Pseudophilautus macropus* + *P. sordidus*, *P. fergusopnianus + P. sarasinorum*]. Leaf litter forms arose five times from [*P. caeruleus*, *P. tanu*, *P. mittermeieri*, *P. zorro* + *P. cuspis* and *P. simba* + *P. semiruber*] common ancestor. The Generalist form may have evolved three times from the common ancestor of [*P. silus* + *P. microtympanum + P. lunatus*]. Canopy forms, too, seem to have evolved at least three times from the common ancestor of [*P. reticulatus* + *P. stellatus* + *P. cavirostris*]. It is of importance that Rock-boulder, Leaf-litter, Generalists and Canopy forms always evolved from a Tree-shrub ancestor and did so during Mid-Miocene to early Pliocene.

**Fig. 4.**
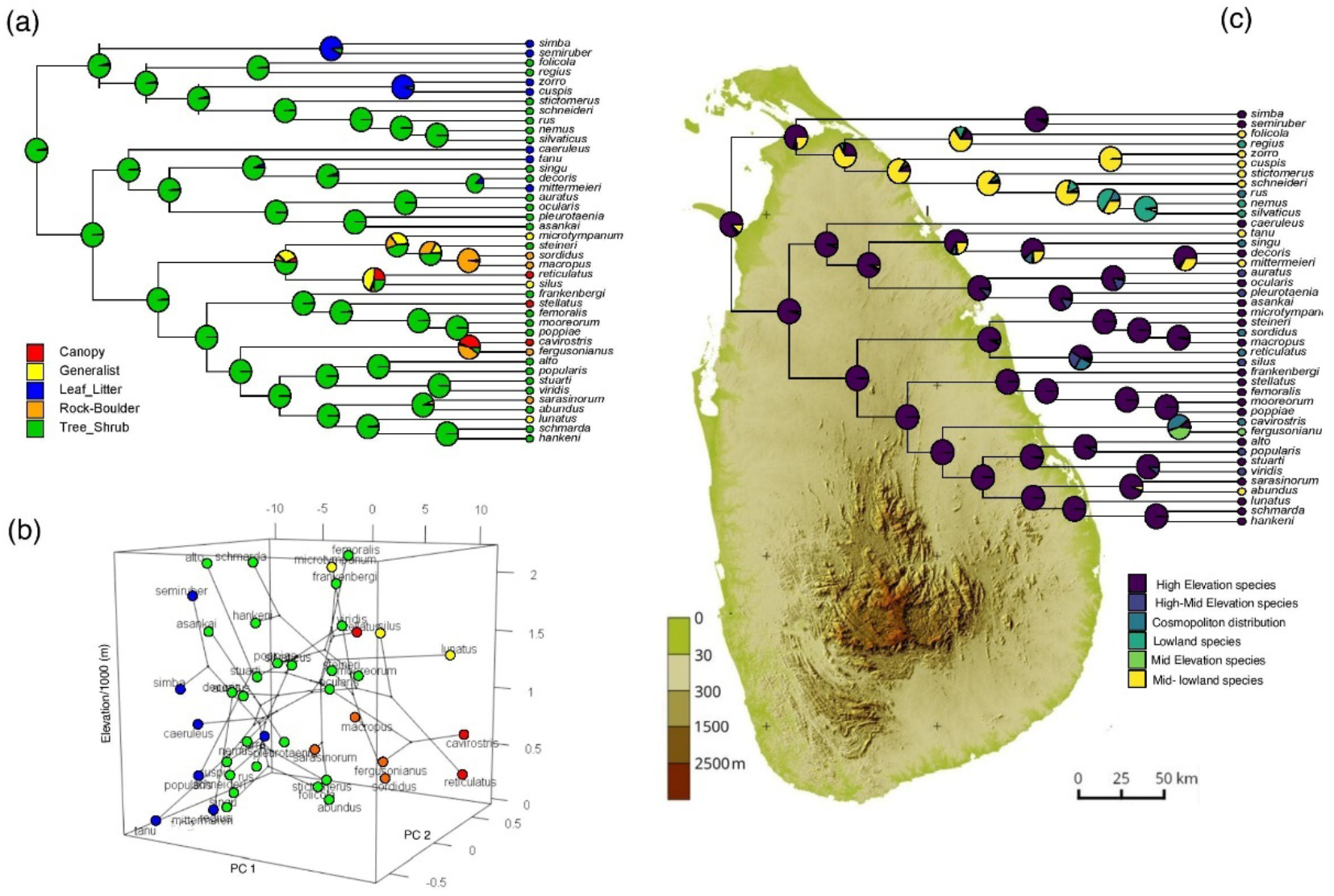
Ancestral state reconstructions (ANC) and phyloecospace of Sri Lankan *Pseudophilautus*. (a) ANC based on five ecomorph categories of *Pseudophilautus*. (b) Phyloecospace of ecomorph categories showing distribution of ecomorphs at different elevations. (c) ANC based on the distribution of species at different elevations. Map of Sri Lanka in the background depict different elevational ranges: highlands—above 1500 m, mid-elevations—500 to 1500 m, and lowlands—below 500 m. The likely ancestor of *Pseudophilautus* is a Tree-shrub form that inhabited the highlands. Elevation subsequently played a role in the evolution of different ecomorphs. For example, Generalists are found only at high elevations while Rock-boulder forms are found at only mid- to low elevations.

The 3D phyoecospace (Fig. 4b) suggests that Tree-shrub forms and Leaf-litter forms are distributed at all elevational levels (highlands—above 1500 m, mid elevations— 500 to 1500 m, and lowlands—below 500 m, Fig. 4c). Rock-boulder forms and Canopy forms are found at mid-to-low elevations, whereas Generalists are found only at high elevations.

### Climatic specializations

The 3D PCA plot depicts the climatic niche occupation in *Pseudophilautus* (Figs. 5a, Extended Data Figure 5) and illustrates how the five ecomorphs occupy different climatic niches, but there is a lot of overlap between them. The first two principal components explain about 88% of the variance (Table S5; Extended Data Figure 5) with PC1 mostly influenced by seasonality in rainfall (bio16, bio17, bio18, and bio19), temperature stability (bio1, bio2, and bio3), and elevation. Tree-shrub forms have expanded their niche along all axes, while Canopy forms and Leaf-litter forms occupy a wider niche along PC1, and Rock-boulder forms and Generalists have expanded niches along PC2. PC2 is mostly a temperature component, dominated by temperature annual range (bio7), which explains about 17.2% of the variance. The third PC, which explains about 6.5% of the variance, also represents environmental temperature (dominated by mean temperature of coldest quarter, bio11, and mean temperature of coldest month, bio6). In the PCA plot, higher PC values correspond to cool and wet conditions, whereas lower PC values indicate warm and dry conditions. The climate niche space of *Pseudophilautus* seem to be determined mainly by temperature variation than rainfall.

During the early to mid-Miocene, approximately 20 to 12 million years ago, the climatic niche space of ancestral Tree-shrub forms of *Pseudophilautus* expanded (Extended Data Figure 5). However, climatic niche evolution along PC1 was gradual and does not follow a clear pattern among different ecomorphs (Fig. 5b, Extended Data Figure 5; PC1. This suggests that ecomorphs evolved through adaptation to local habitat conditions rather than specific macroclimatic conditions. Nevertheless, deviations from ancestral climatic conditions were minimal along PC2 (Extended Data Figure 5; PC2) and PC3 (Extended Data Figure 5; PC3). Rock-boulder forms evolved towards warmer and drier conditions, with species distributions affected by ranges of extreme temperature conditions and less cold tolerance. Niche expansion along PC2 and PC3 was quite recent for *Pseudophilautus*, occuring around the late Miocene and early Ploicene periods (Extended Data Figure 5; PC2 and PC3). The disparity-through-time plots suggest that the rate of evolution of climate niches increased during the second quarter of the evolution along PC2 (Fig. 5c) and during the fourth quarter of along PC3 (Fig. 5d). These intervals correspond to the mid Miocene and late Pliocene periods, when conditions became dry and warm during the mid Miocene glaciation (Mi-1 glaciation) and wet and cold when Asian monsoons intensified during early Pliocene respectively. However, the rapid expansion of climate niches along all axes seems recent in *Pseudophilautus*.

**Fig. 5.**
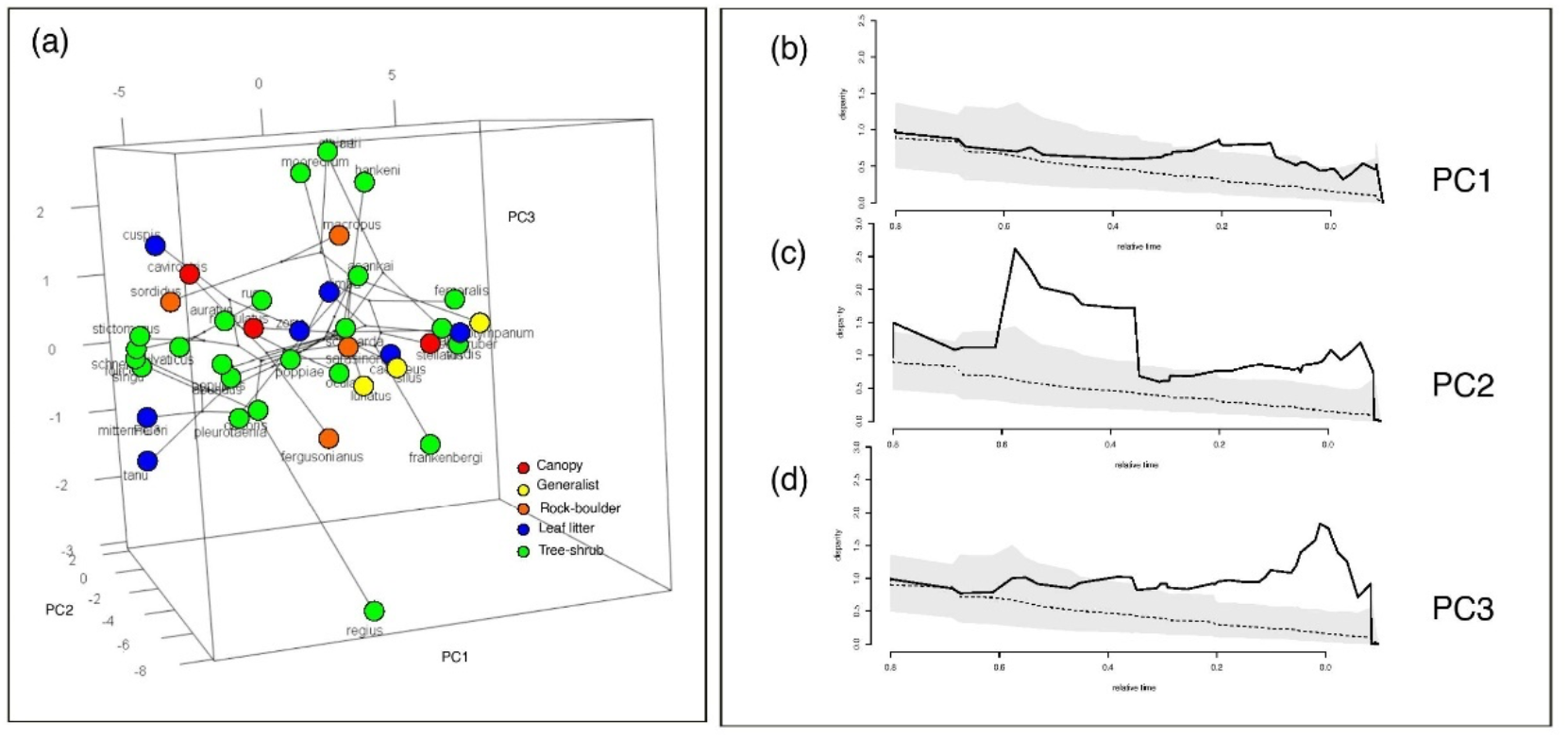
Climate-niche occupation of *Pseudophilautus* ecomorphs in 3D phyloecospace and the evolution of climate niches of different ecomorphs. Although a dominant pattern is not evident in the occupation of the the climatic niche space, Leaf-litter forms appear to have a wider niche breadth along PC1 and generalists and canopy forms have an expanded niche along PC2. Relative disparity-through-time (DTT) plots of PC scores: PC1 (b), PC2 (c) and PC3 (d) depict change in the rate of climatic niche evolution. Solid lines indicate the observed DTT; dashed lines and corresponding polygons represent averages and 95% confidence intervals of the expectations given a constant accumulation of disparity over time based on 999 pseudo-replicates. A higher rate of climatic niche evolution is observed during the second and fourth quarters of the evolutionary history of *Pseudophilautus*. These intervals correspond to mid-Miocene climatic cinditions and monsoon-driven, wet Pliocene climatic conditions, respectively.

## Discussion

The adaptive radiation of *Pseudophilautus* in Sri Lanka is characterized by an early burst radiation, high species diversity and common ancestry, the typical features of recognized adaptive radiations[3,5,6,36,37](Glor 2010; Losos and Ricklefs, 2009; Losos & Mahler 2010; Schluter 2000; Givnish 1997). While strict monophyly of a clade is not always required to be classified an adaptive radiation, recognizing unusual lineages can help identify the processes that limit morphological disparification, species richness, and unusual distributions [38-42](Brandley & De Queiroz, 2004; Johnson et al. 1993; Clarke et al. 1996; Jackman et al. 1999; Rodriguez-Robles et al. 2007). The Indian mainland clade of *Pseudophilautus*, comprising only three species, does not exhibit ecomorph resolution, which suggests that a lack of ecological opportunity in the mainland may have hindered ecological diversification in this clade. In contrast, the island radiation of Sri Lankan *Pseudophilautus* shows a clear morphology-habitat correlation, with recently evolved ecomorphs. Furthermore, the initial colonization from the mainland offered an ecological opportunity for Sri Lankan *Pseudophilautus*. This led to their rapid diversification, as evidenced by the negative gamma value in lineage plots (Fig. 2), during the early phases, and subsequently slowed as ecological niches were occupied.

The ecomorph diversification of the extant *Pseudophilautus* radiation can be traced back to the late Miocene. This timeline implies that following its initial colonization, a habitat-specialist ancestral species underwent swift diversification, leading to the emergence of ancestral ecomorph types. This initial diversification is marked by a notable shift towards larger body sizes in these ancestors. However, these early ecomorphs eventually faced extinction. Consequently, by the late Miocene, as environmental conditions turned favorable, these niches were occupied by the extant ecomorphs.

Analysis of morphology reveals distinct morphological groupings (i.e., ecomorphs) associated with each of the habitat types we postulate (Fig 1, Extended Data Figure 1). *Pseudophilautus*, the largest of which are tree canopy forms, require larger hands and feet, and wider disk widths to jump over larger distances. The smallest frogs are leaf litter forms, but there are two distinct types of morphology within this group [43](Bahir et al. 2005). The diminutive forms (less than 15 mm SVL) have short eye-snout distances and almost no disks, while slightly larger frogs (*P. cuspis* and *P. zorro*) have proportionately the largest eye-snout distances and moderately developed disks with flattened bodies, which may aid movement in leaf-litter [29](Manamendra-Arachchi and Pethiyagoda, 2005). The diminutive forms are small enough to move around in leaf litter, while the Generalists also inhabit this habitat [31](Meegaskumbura et al. 2012). Rock-boulder forms have greater head widths and higher internarial and interorbital distances, resulting in lateral widening. These observations affirm that the morphology of *Pseudophilautus* species is highly correlated with specific habitat types. Although, the clades of closely related species are typically composed of similar ecomorphs, there are several instances where ecomorphs have evolved independently from different lineages (Extended Data Figure 1).

Accelerated phenotypic divergence at the initial stages is a common feature of most adaptive radiations [4,18,20](Losos and Miles, 2002; Erwin, 1994; Simpson, 1953). Within the *Pseudophilautus* radiation, alongside the rapid diversification of various phenotypes, there’s a clear surge in early speciation rates, underscoring its adaptive nature. Although earlier ecomorphs appear to have faced extinction, they subsequently paved the way for the emergence of new ecomorphs. This is evident by the ecomorphs (Leaf litter, Rock-boulder and Generalists) deriving from the shrub-tree ecomorph arise only well into the diversification event during the late Miocene, Pliocene and early Pleistocene. The bayou analysis confirms an initial shift towards larger body sizes followed by later shifts towards decreasing body sizes (Fig 3b). This supports a dynamic fitness landscape for *Pseudophilautus* where, during some period of evolution, larger body sizes were selected over smaller; and later, under favorable environmental conditions, smaller body sizes came to be selected for. These changes may have been driven by environmental changes during different periods (e.g.: change in vegetation from broad leaved forests to smaller leaves, extreme climate changes driving conserved body plans for heat conservation). Furthermore, larger body sizes may also have exerted negative selection pressures such as increased predation risk during early evolution such that smaller body sizes were favored later on. However, there is insufficient evidence at present, to validate these arguments.

The ancestral reconstructions of body size suggest that the adaptive radiation of *Pseudophilautus* species may have originated from a small frog that was likely a shrub dweller (Fig. 3a). As the radiation progressed, there appears to have been an initial burst of diversification to fill micro-niches within the shrubs and eventually into the canopy. However, as the available niches became saturated, competition may have driven adaptive diversification into other ecomorph types, such as the Leaf litter, Rock-ground, and Generalists. This raises an important evolutionary question about the role of interspecific competition in driving phenotypic diversification [44](Svanbäck and Bolnick, 2007).

During the early stages of radiation, ecological opportunities would be plentiful due to the existence of unoccupied niches [45](Day and Young 2004). However, the likelihood of adaptation to new niches may have decreased if divergent natural selection in the context of competition was weak [46-48](Gavrilets and Vose, 2005; Dayan and Simberloff, 2005, Gavrilets and Losos, 2009). As the empty niches became filled, competition among shrub-tree dwellers increased, potentially leading to diversification into other habitats such as leaf litter (which favored smaller-sized morphs) or rock-boulders (which favored larger-sized species better equipped to compete with near-water typical amphibians). In addition, larger-bodied species may have experienced negative selection pressure due to predation, which could have restricted their diversification and resulted in many smaller species and fewer larger species.

Ancestral-state reconstructions of the root node of *Pseudophilautus* (Fig. 4a and 4c) suggest that the ancestor was a Tree-shrub species from high elevations, specifically the central-hills of the island [28](Meegaskumbura et al. 2019). Three of the ecomorph types, Tree-shrub, Leaf litter and Rock-boulder, are present in both lowlands and highlands, but the Generalists occur only at higher elevation (above ∼700m asl). The Rock-boulder morphs are found along boulder-strewn streams from the highlands to the lowlands; the Leaf litter forms are found in canopy-covered forests of both high and low elevation forests but never in open habitats that lack leaf litter. Tree shrub morphs show the highest distribution, occurring in both open and closed habitats of the highlands and the lowlands. Contrastingly, the Generalists, which occupy all three habitat types, are found only in the cooler and wetter higher elevations (Fig.4b) [29](Manamendra-Arachchi and Pethiyagoda, 2005).

Since the ancestral-state reconstructions of the root node of *Pseudophilautus* suggests that the ancestor was from high elevations, specifically the central-hills of the island [28](Meegaskumbura et al., 2019), it can be presumed that it was a cool-and-wet-adapted, Tree-shrub-dwelling specialist that seeded the species pump on the island. The Generalists seems to have been derived secondarily; Generalists, in the context of the Sri Lankan *Pseudophilautus* radiation, are species that occupy several habitat types (i.e., Tree-shrub, Rock-boulder and Leaf-litter). In *Pseudophilautus*, the few Generalist species are nested within two disparate clades (Fig. 2). This pattern in habitat utilization of generalists bolsters the notion that terrestrial direct development expresses its full potential in cool and humid habitats. Supporting this hypothesis, the Generalist clade is distributed at high elevations (over 700 m asl) in the perennially wet cloud forests, grasslands, and associated habitats of the three main mountain ranges. It is likely that only in these cool, wet habitats that direct developing species could evolve a more expansive niche breadth, because of their reliance on high humidity for breeding and egg development [25,43,49](Meegaskumbura et al. 2015; Bahir et al. 2005; Schweiger et al. 2017).

The late Miocene coincides with the evolution of the climatic niche of *Pseudophilautus* into extreme temperature conditions [50,51](Chatterjee et al. 2013; Ellepola et al. 2022) along PC2 (Fig. 5c), during which the evolution of ecomorphs preceded. However, by the late Miocene, Himalayan-Tibetan orogeny, which started around 30 MYA was well advanced, paving way for two stable monsoons per year [50,52-54](Chatterjee et al. 2013; Yin & Harrison, 2000; Gehrels et al. 2003, Ellepola and Meegaskumbura, 2023). This in turn aided in the evolution of more ecomorphs during the early Pliocene. Further morphological disparification (formation of other ecomorphs such as Leaf litter, Canopy, Rock-boulder and Generalists) of *Pseudophilautus* seems to have happened during the late Miocene or Pliocene, with the creation and intensification of the two monsoons and lowering of the temperature in the region (Figs. 4 and 5).

Our analyses sheds light on the evolution of ecomorphs in the *Pseudophilautus* radiation of frogs and suggest that ecomorphs evolved multiple times across the phylogeny, in different geographic regions, and at different altitudes, primarily during a cold climatic period, possibly in association with monsoon cycles arising from Himalayan-Tibetan orogeny. The common ancestor of the radiation was possibly a medium-sized, wet-adapted, tree-shrub-dwelling habitat specialist, and extreme size classes evolved relatively recently. We highlight the importance of considering ecomorphs in understanding the adaptive radiations of frogs and suggest that adaptive radiations, subtler than in tailed vertebrates, may be more prevalent in frogs than previously recognized. Our findings add to the growing body of knowledge on the ecomorphology and adaptive radiations of frogs and underscore the need for further research.

## Methods

The genus *Pseudophilautus* in Sri Lanka represent a spectacular diversification comprised of direct developing frogs. Seventy-two species are recognized at present [21](AmphibiaWeb, 2023) out of which three (*P*. *wynaadensis*, *P*. *amboli* and *P*. *kani*) represent a subclade from India. The Sri Lankan diversification comprise of 16 extinct species [33](Meegaskumbura et al. 2007) as well as 10 nominal species lacking genetic data [55](Ellepola et al. 2021). We included 41 species of Sri Lankan *Pseudophilautus*, whose validity has been confirmed based on the general lineage concept [56](de Quiroz, 1999), in the current analysis. These species occupy several habitat types ranging from open grasslands, rock substrates along streams, rainforest canopy, shrubs, leaf litter and anthropogenic habitats, and most are confined to higher elevations of the wet, montane region of the island’s southwestern quarter [28,29,33](Manamendra-Arachchi and Pethiyagoda, 2005; Meegaskumbura et al., 2007, Meegaskumbura et al. 2019). To serve as the outgroup, two species from the Indian sister group, *R*. *charius* and *R*. *signatus* were included in the phylogenetic analyses.

### Morphometrics

Thirty-two morphological measurements from adults of both sexes of all species were obtained following the terminology of [29]Manamendra-Arachchi & Pethiyagoda (2005). All measurements were made by the same person to avoid measurement bias (Data S1). Abbreviations: distance between back of eyes (DBE), distance between front of eyes (DFE), length of disk of third finger (DL), width of disk of third finger (DW), eye diameter (ED), eye-to-nostril distance (EN), eye-to-snout distance (ES), femur length (FEL), finger length (FL), foot length (FOL), head length (HL), head width (HW), inner metatarsal tubercle length (IML), internarial distance (IN), interorbital width (IO), lower arm length (LAL), mandible–back of eye distance (distance from angle of jaws to posterior-most point of eye; MBE), mandible–front of eye distance (distance from angle of jaws to anterior-most point of eye; MFE), mandible–nostril distance (distance from angle of jaws to middle of nostril; MN), nostril–snout distance (distance from middle of nostril to tip of snout; NS), palm length (distance from posterior-most margin of inner palmar tubercle to tip of disk of third finger; PAL), snout-vent length (SVL), tibia length (TBL), toe length (TL), tympanum to eye diameter (TYE), tympanum diameter (TYD), upper arm length (UAL) and upper eyelid width (UEW). Digit number is represented by roman numerals I-V (Data S1).

### Morphology-habitat correlation

Five main habitat type occupations have been proposed for Sri Lankan *Pseudophilautus* [29-33](Manamendra-Arachchi and Pethiyagoda 2005; Meegaskumbura and Manamendra-Arachchi 2005, 2007, 2009, 2012): Canopy, Generalist, Rock-boulder, Leaf litter and Tree-shrub. These may represent ecomorphs.

Mean values for all mensural data were calculated for each sex of each species and were log10-transformed to fulfill assumptions of normality and homoscedasticity. To explore morphological variation among species, multivariate principal component analyses (PCA) were performed using habitat as the grouping variable. Despite the sexual dimorphism evident in *Pseudophilautus* species [29,30](Manamendra-Arachchi & Pethiyagoda 2005, Meegaskumbura & Manamendra-Arachchi 2005), adult males and females were included in the same analyses but identified by sex in the resulting plots. However, because body size may obscure shape variation in interspecific morphological data, we performed a PCA based on size-corrected data as well [57](Velasco and Herrel, 2007). This enabled us to distinguish between size and shape of the species being analysed. To remove effects of body size, residuals were calculated of the regressions of the log10-transformed morphological variables against log10-transformed SVL. The residual values were then used in a new principal component analysis, to explore variation in shape among the species. Additionally, to account for the phylogenetic non-independence among species, phylogenetic principal component analyses (pPCA) were also performed on the data sets.

Linear discriminant analysis (LDA) was performed on normalized measurement data as well as on residual values (i.e., the size-corrected data) to predict the probability of a species belonging to an ecomorph category based on multiple predictor variables. Discriminant analysis maximizes the differences between groups as opposed to maximizing variance along axes in a PCA. The LDA determines group means and computes, for each species, the probability of belonging to a given ecomorph category. The species is then allocated to the category with the highest probability score. Here, 60% of the data were treated as training data and 40% were treated as testing data in the LDA model.

### Phylogenetic analysis

To infer relationships among *Pseudophilautus*, we constructed a multi-gene phylogeny based on three mtDNA (16S rRNA, 12S rRNA and CytB) and the NucDNA Rag-1 molecular loci. The data for CytB gene fragments were newly generated during the current study (others obtained from [28]Meegaskumbura et al. 2019), and for this, DNA was extracted from ethanol preserved tissues using Qiagen tissue extraction kits following manufacturer’s protocols. Portions of the CytB genes were amplified by PCR and sequenced directly using dye-termination cycle sequencing. Thermocycling conditions included an initial step at 94 °C for 3 min, followed by 35 cycles at 45 s at 94 °C, 1 min at 46–50 °C and 45 s at 48–56 °C, and a final step at 72 °C for 5 min. GenBank accessions used in the analyses are listed in Table S5.

Mitochondrial gene fragments were aligned using MUSCLE as implemented by MEGA v.11 [58](Tamura et al. 2021); regions for which there was low confidence in positional homology were removed from the analysis. Nuclear gene sequences were aligned using MEGA v.11 with translated amino-acid sequences. The complete concatenated dataset included 43 valid taxa [55](Ellepola et al., 2021) and a total of 2702 base pairs (bp). Two species of *Raorchestes* (*R. charius* and *R. signatus*), the sister group of *Pseudophilautus*, were included as outgroup taxa.

Tree topology was inferred both using a maximum likelihood approach (IQ-TREE; [59]Nguyen et al. 2015) and a bayesian approach (BEAST v.2.5; [60]Drummond and Rambaut, 2007) of which both yielded similar results, hence the tree topology obtained from BEAST were used in subsequent analyses. Both analyses were performed for both partitioned and unpartitioned datasets as well as for each gene individually. The dataset was partitioned into specific gene regions by specifying character sets (charset 12S rRNA = 1–308; charset 16S rRNA = 309–776; charset CytB = 777–1310; charset Rag-1 = 1311-2702). The partitioned dataset was used for the phylogenetic analyses. The best-fitting nucleotide substitution model for each dataset was chosen using jModelTest v.2.1.4 [61](Dariba et al. 2012). Model GTR+I+G as the nucleotide substitution model, Yule model as the tree prior, and lognormal relaxed clock as the molecular clock were assigned in BEAST and the analysis was run on the Cipres Science Gateway Server [62](Miller et al. 2015) for 50 million generations in two consecutive runs. Burnin was defined by observing the log-output file in Tracer v.1.6 [63](Drummond et al. 2012); 90% of post-burnin trees were analysed using Tree Annotator and a final maximum clade credibility tree constructed (Data S2).

To examine the temporal context of divergence, we estimated divergence times among lineages on the partitioned dataset by using BEAST v.2.5 [60](Drummond and Rambaut, 2007). The Yule model was given as the tree prior, the GTR+I+G model of evolution specified, and other parameters estimated for all gene partitions. Over 10 million generations of Markov chain Monte Carlo (MCMC) simulations were run, with one tree saved per 1000 generations. The analysis was run using the uncorrelated lognormal relaxed clock. We calibrated the tree using a literature-based, molecular-estimated secondary calibration point from [64](Chen et al. (2020) (i.e., the age of the most recent common ancestor (MRCA) of extant *Pseudophilautus*, 16.48–24.92 Mya) which was derived based on different calibration prior-combinations that included fossil calibrations as well as secondary calibrations on a Anchored Hybrid Enrichment data set.

We analysed the accumulation of lineages through time to discern the tempo of diversification in the radiation [65](Harmon et al., 2008). Lineage-through-time plots were constructed by plotting the log number of lineages against the divergence time as implemented in phytools 2.0 [66](Revell, 2023). A null model was also developed wherein the tree was simulated with a total of 72 taxa (to account for the missing taxa-[21]Amphibiaweb (2023) records 72 species for *Pseudophilautus*) and assuming a constant rate of speciation across the lineages (under a pure birth model) with no extinctions. A Monte Carlo Constant Rate test (MCCR) was used to calculate the gamma (γ) statistic in phytools 2.0, which compares the general shape of the lineage through time curve to a null pure birth model. If γ < 0, the radiation shows an early, rapid speciation event; if γ = 0, the radiation undergoes a constant diversification rate; and if γ > 0, the diversification rate increases towards the present [67](Pybus and Harvey, 2000).

We visualized the evolution of traits of *Pseudophilautus* defined by PC1 and PC2 axes in a phylomorphospace [68](Sidlauskas, 2008) using the phylomorphospace function in PHYTOOLS [69](Revell, 2012). Further, by using principal component values, we tested whether there are significant differences among ecomorphs in their morphospace positions by performing a phylogenetic MANOVA. This was backed by phylogenetic ANOVA and post hoc tests performed on each PC axis treated with ‘ecomorph’ as the categorical variable [69](Revell, 2012).

Morphological variation in *Pseudophilautus* was described by estimating ancestral states of PC1 and PC2 and visualizing them using traitgrams [70](Evans et al. 2009) under a browninan motion model, implemented with the phenogram function in PHYTOOLS [69](Revell, 2012). Ancestral states were reconstructed separately based on different ecomorphs, and elevational level (primarily: highlands—above 1500 m, mid-elevations—500 to 1500 m, and lowlands—below 500 m; species sharing two or more levels were categorized as high-mid, mid-low or cosmopolitan; Data S3). To infer the number and timing of evolutionary shifts within ecomorphs, we used stochastic character mapping [71,72](Neilson 2002; Heulsenbeck et al. 2003) implemented with the make.simmap (model=“ER”) function in PHYTOOLS [69](Revell, 2012). We tested multiple models for the ancestral state reconstruction; the ER model gave the highest likelihood. On each of the 1000 post burnin trees obtained from BEAST, we used stochastic character mapping to generate 100 potential histories. This approach considers uncertainty in both the evolutionary history of the traits as well as the inferred topology of the phylogeny.

To identify evolutionary shifts in body size (defined by PC1 axis), the bayou v. 2.1.1 analysis [73](Uyeda and Harmon 2014) was implemented in R v. 4.2.1 [74](R Core Team 2022). The ‘bayou’ approach detects evolutionary shifts towards different optima without influence by a priori groupings on the tree. Bayou uses a reversible-jump Bayesian approach that uses univariate data to estimate the placement and magnitude of evolutionary shifts [73](Uyeda and Harmon 2014). We used an unconstrained bayou analysis on PC1 values using an OU model with different priors. The model having the prior values alpha = 0.09, sig2 = 0.09, k = 0.25, theta = 0.32, and loc = 0.01 was selected as a reasonable OU model for the analysis, and shifts with a posterior probability (pp) of 0.5 were regarded as significant. We used PC1 values, which explain 93.2% of the variance, as the continuous variable, as it mostly represents body size.

### Climatic niche evolution

We obtained from published literature present-day geographical occurrences in the form of GPS coordinates for all extant *Pseudophilautus* species, which we supplemented with geocoordinates from our own field data. Location data are provided in Data S3. Our final data set includes a total of 363 occurrence records.

Information on 19 bioclimatic variables and elevation for each occurrence point (at a spatial resolution of 0.5 arcsec) was obtained from WORLDCLIM 2.0 [75](Ficks and Hijmans 2017) using the “extract” function in RASTER 3.0-7 [76](Hijmans et al. 2015). Mean values for each bioclimatic variable for each taxon are provided in Data S4.

We used the average bioclimatic conditions in a PCA based on their correlation matrix to illustrate climatic niche occupation of different ecomorphs. The axes to be retained for further analyses were determined using the broken-stick method as implemented in VEGAN 2.5-6 [77](Oksanen et al. 2017). We assumed that measured species means are a reasonable approximation of the realized climatic niche of the species [78](Wiens et al. 2007). We constructed the 3D phyloecospace for all ecomorphs by using the calculated PC scores to show the climatic niche occupation of ecomorphs. We then assessed the extent to which traits (climatic niche evolution along each PC axis) had accumulated over time in each biogeographic region using disparity-through-time (DTT) plots [79](Harmon et al. 2003), with expected disparities calculated based on 1000 resampling by using the ‘dtt’ function in GEIGER 2.0.6.2 [65](Harmon et al. 2008) and with phenograms (projections of the phylogenetic tree in a space defined by phenotype/PC axis and time) constructed with the ‘phenogram’ function in PHYTOOLS [69](Revell, 2012).

## Data availability

The authors declare that data supporting the findings of this study are available within the Data files S1, S2, S3 and S4. Additional data that support the findings of this study are publicly available online at https://www.ncbi.nlm.nih.gov/genbank/, http://www.iucnredlist.org/technical-documents/spatial-data, https://www.gbif.org and https://www.worldclim.org. For Genbank accession numbers see Table S5.

## Code availability

All codes used in support of this publication are publicly available in specific R packages which we have mentioned in the reference list.

## Acknowledgements

We thank the Department of Wildlife Conservation and the Forest Department of Sri Lanka for permission to carry out rhacophorid-based research in Sri Lanka. Wildlife Heritage Trust of Sri Lanka (RP) and the National Museum of Sri Lanka has provided access to specimens and facilitated our amphibian research over several decades.

## Funding

This work was supported by the Wildlife Heritage Trust of Sri Lanka, US-NSF funding to (CS), Ziff Environmental Postdoctoral Fellowship, Harvard University Center for the Environment (MM), Guangxi University Laboratory Startup Funding (MM) and China Student Council Fellowship for graduate studies (GE).

## Author contributions

Conceptualization, M.M., M.P., G.E. and G.S.; methodology, M.P. and G.E.; formal analysis, M.M. and G.S, G.E,. M., N.W. and MP; investigation, M.M.; data curation, G.E., G.S., D.S; writing—original draft preparation, M.M., G.E., G.S.,K.M., N.W. M.P., D.S., R.P. and C.S; writing—review and editing, M.M., G.E., G.S.,K.M., N.W. M.P., D.S., R.P. and C.S; supervision, M.M, R.P. and CS.

## Competing interests

The authors declare no competing interests.

---

**Extended Data Figure 1.**
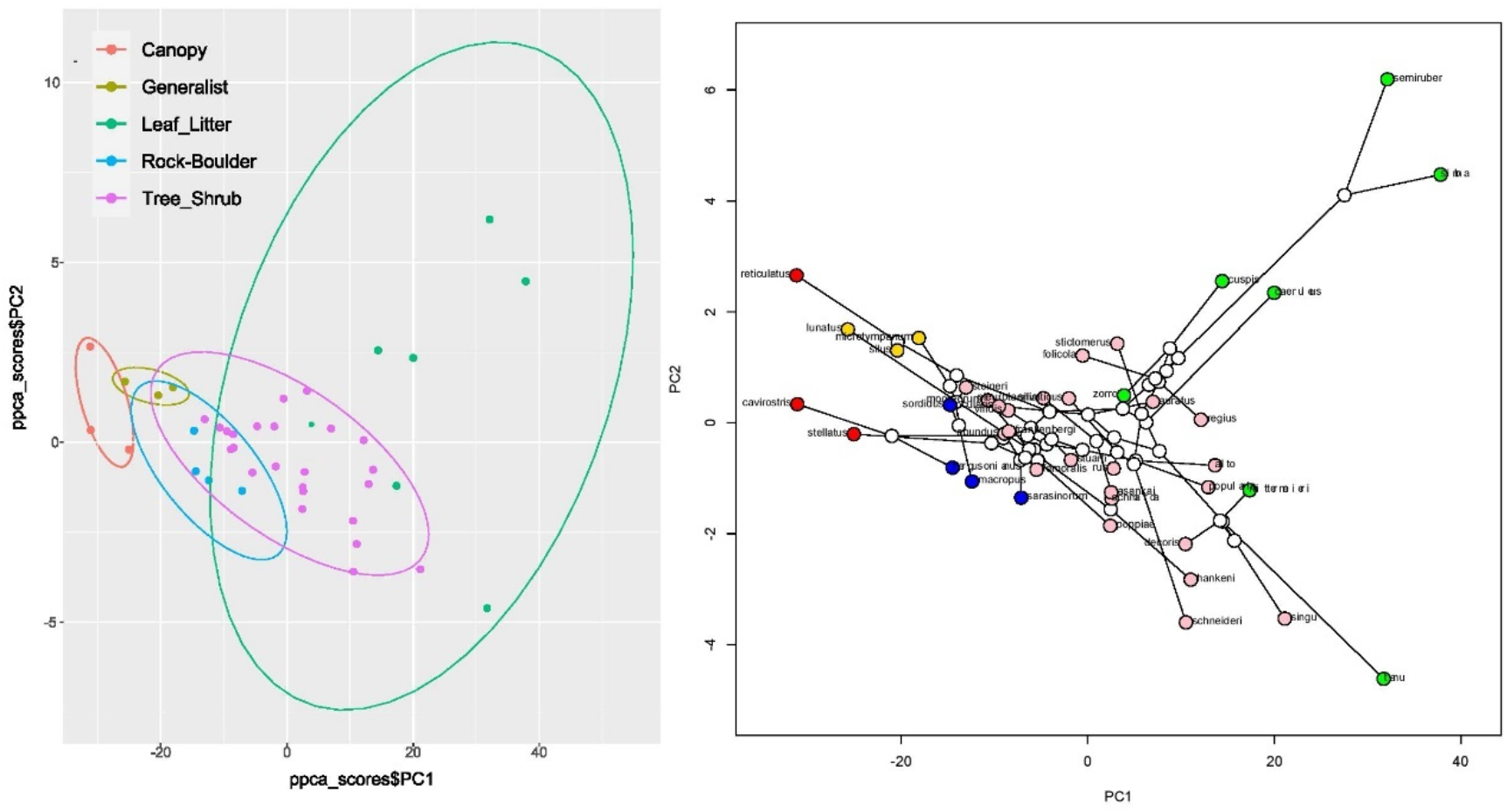
The position of Sri Lankan *Pseudophilautus* in morphological space as indicated by the first two axes of the phylogenetic principal component analysis (pPCA). Ecomorphs categories are distinguished mainly along pPC1 axis (left). pPC1 is primarily defined by body size (SVL), foot length (FOL) and head length (HL) whereas PC2 is mainly defined by upper eyelid width (UEW), lower arm length (LAL) and interorbital width (IN). The phlomorphaspace laid on the pPC axes shows that ecomorphs cluster despite being phylogenetically unrelated hence pointing towards a convergent adaptive radiation in *Pseudophilautus*.

**Extended Data Figure 2.**
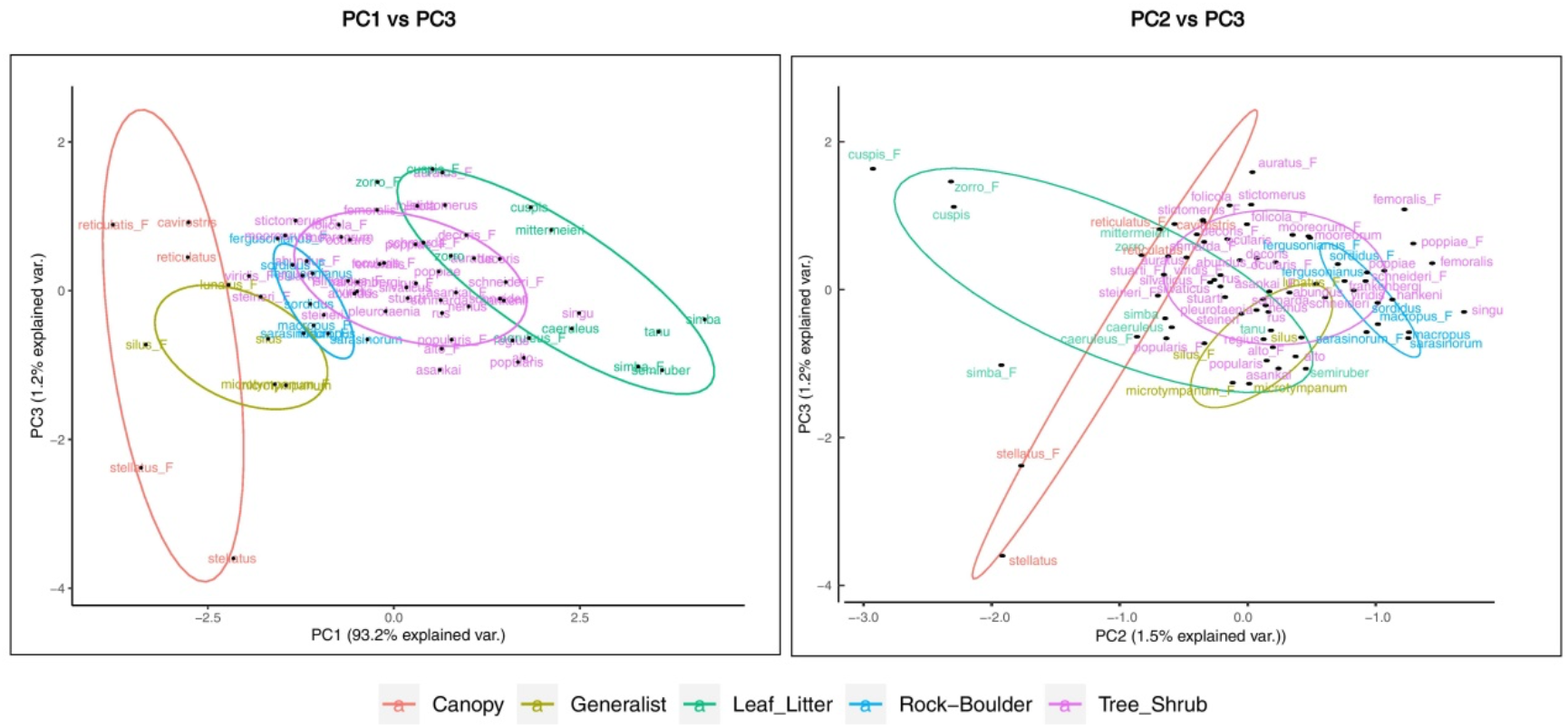
The position of Sri Lankan *Pseudophilautus* in morphological space as indicated by principal component analysis (PC2 and PC3) conducted on the raw data. Ecomorph categories are clearly separated along the PC1 axis. PCA defined by PC2 and PC3 explains only about 2.7% of the variance. PC2 and PC3 do not distinguish between ecomorph categories, although Upper eyelid width (UEW) and Palm length (PAL), and distance from eye to nostril (NE) and Nostril–snout distance (NS), load heavily on PC2 and PC3, respectively (see Table S1).

**Extended Data Figure 3.**
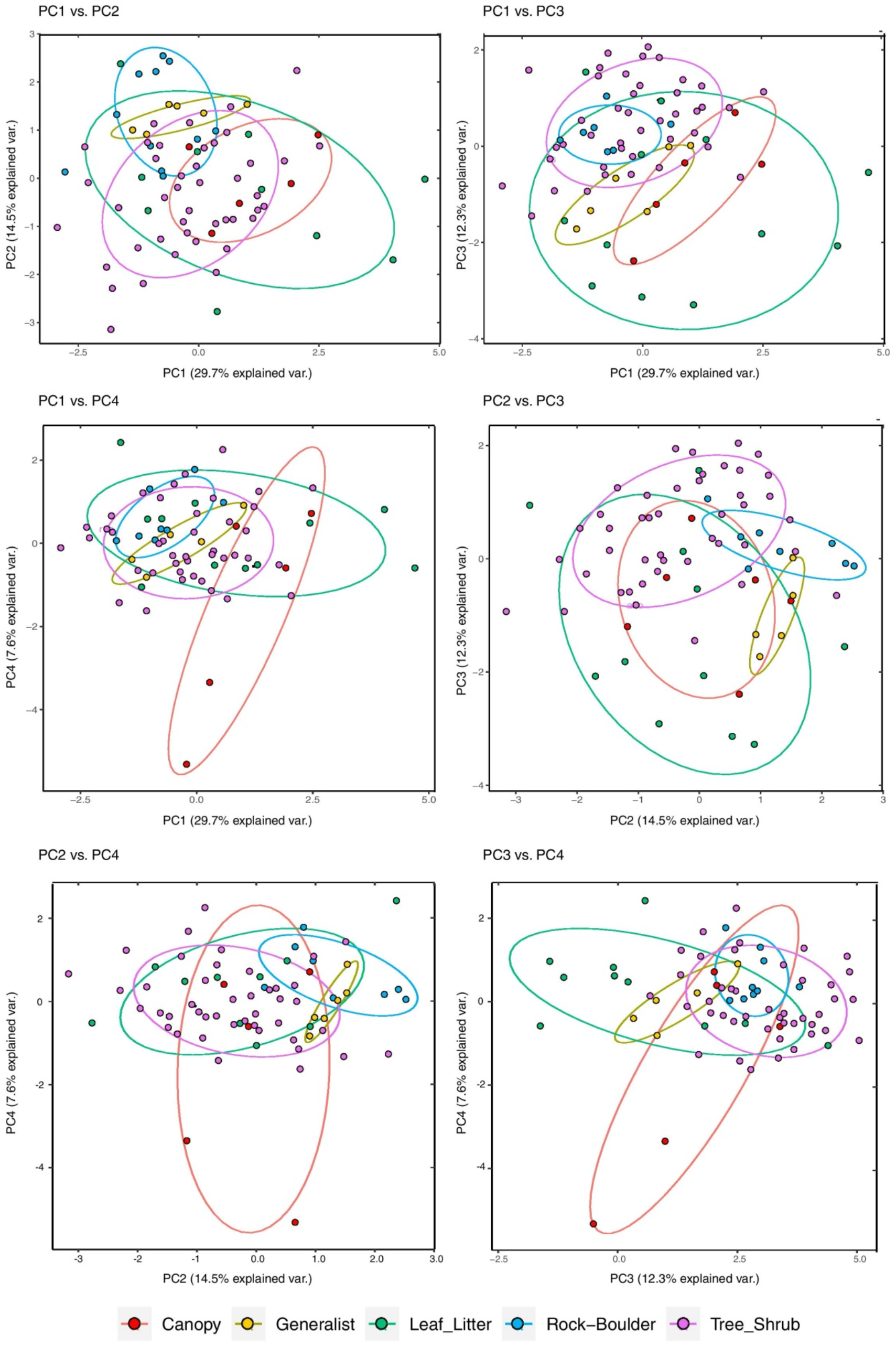
The position of Sri Lankan *Pseudophilautus* in morphological space as indicated by different axis combinations of the principal component analysis, using size-adjusted data. Overall, different ecomorph categories are not well defined along any axis. However, Canopy and Generalist forms segregate along PC2, while Rock-boulder forms segregate from Canopy forms along PC1 and PC3. A general pattern indicates that all other ecomorphs exist within the morphological space of Leaf-litter forms but Tree-shrub forms being an exception along PC4. The relationships between different principal component axes are shown by sub plots.

**Extended Data Figure 4.**
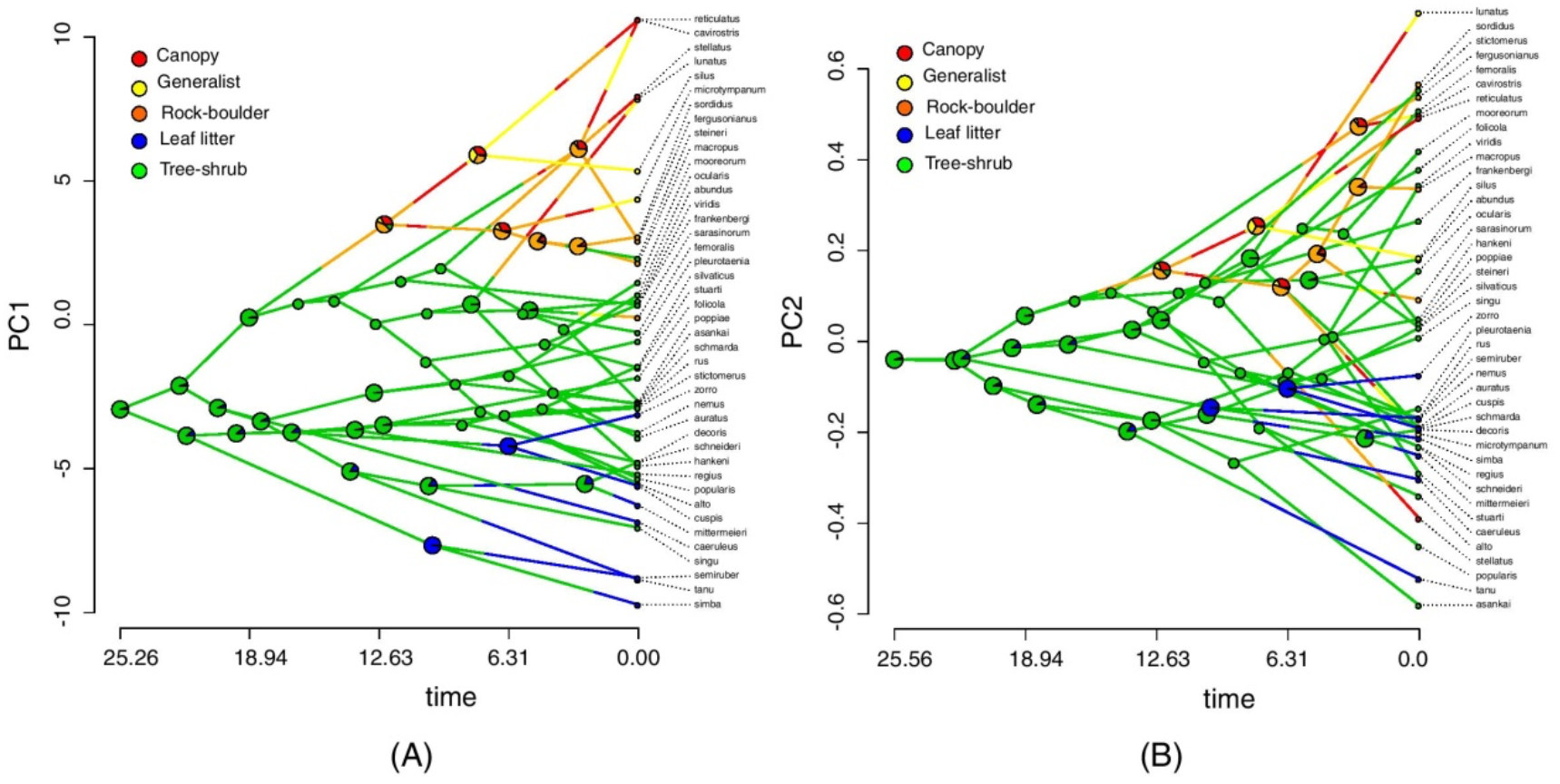
Temporal patterns of morphological evolution in ecomorphs along PC1 and PC2. The traitgram of PC1 (A) suggests that the evolution of different morphs is attained mainly by altering body size, whereas a clear pattern is not evident by the traitgram of PC2 (B). Overall, these patterns suggest that body size evolution and evolution of morphological characters such as Upper eyelid width (UEW) and Palm length (PAL) were largely decoupled during the history of *Pseudophilautus*. Ancestral-state reconstructions traced on traitgrams suggest that a medium-sized tree shrub form has given rise to other ecomorphs. Other ecomorphs likely evolved between 10 and 5 MYA.

**Extended Data Figure 5.**
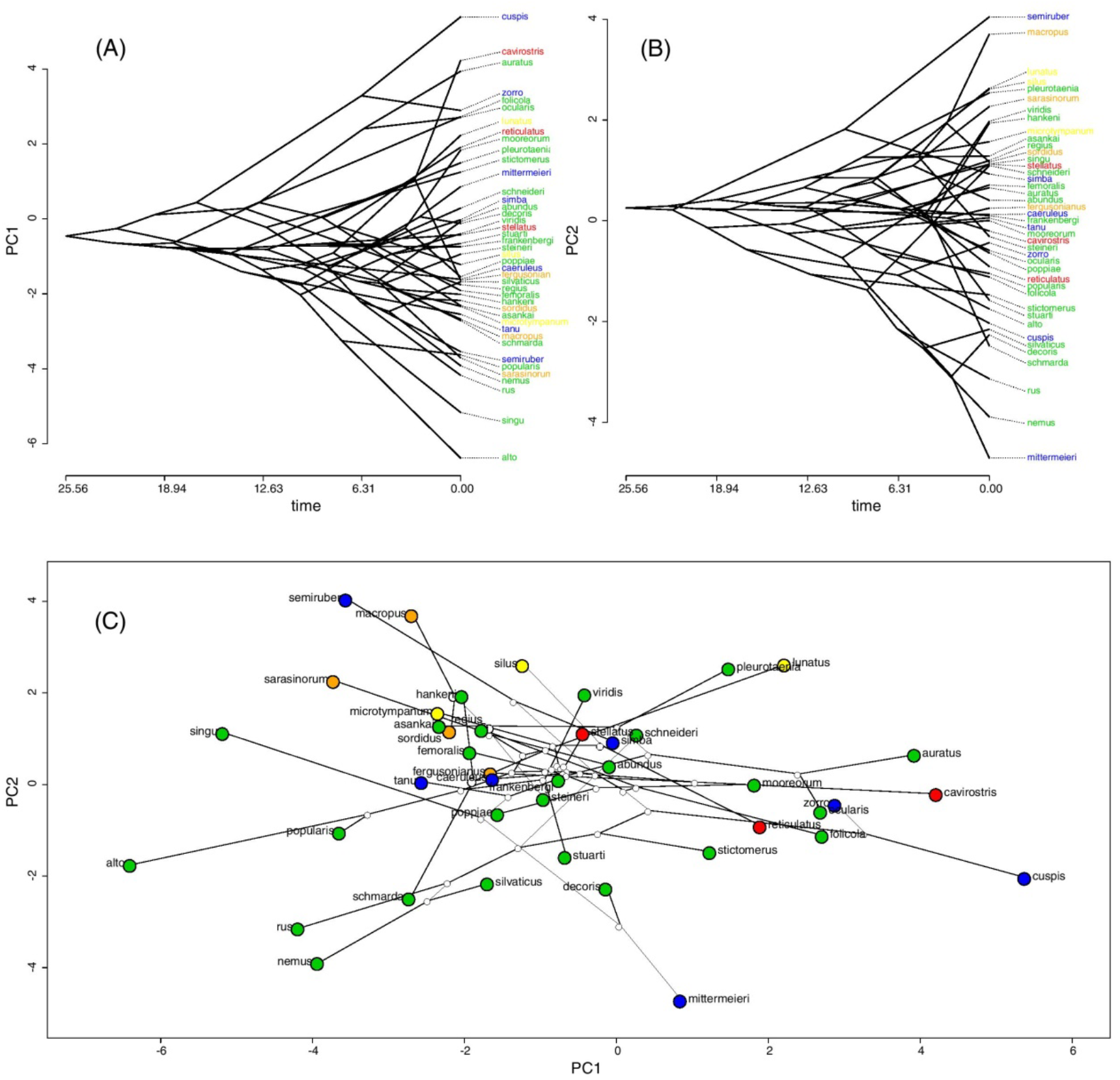
Temporal patterns of morphological evolution along PC1 and PC2, and ecomorphs as phylomorphospace-traitgrams in *Pseudophilautus* (using size-corrected residual data). A clear correlation of body shape and ecomorphs are not evident from the traitgrams. Traitgram of PC1 (A); Traitgram of PC2 (B); Different ecomorphs traced on the phylomorphospace (C) indicating tree shrub forms spanning the entire range of body shapes.

**Extended Data Figure 6.**
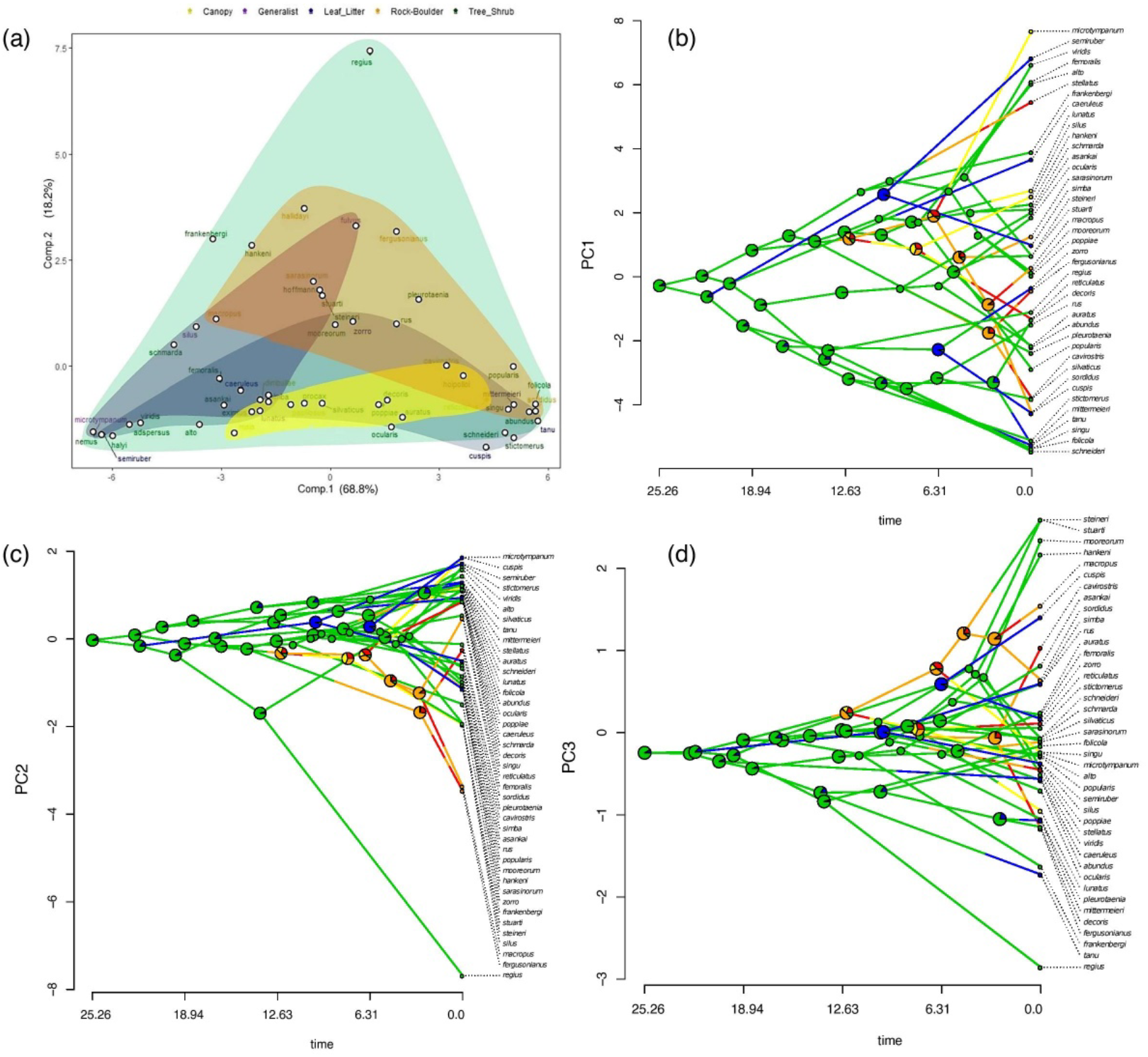
Climatic niche space occupation and climatic niche evolution in *Pseudophilautus* ecomorphs. (a) Climatic niche occupation of Pseudophilautus ecomorphs. Minimum convex **polygons** for different ecomorphs are shown by different colours. Climatic niches of the ecomorphs largely overlap, although there is some separation among Canopy forms, Rock boulder forms and Generalists. Canopy forms and Leaf litter forms appear to have expanded their climatic niche along PC1, whereas Generalists and Rock boulder forms have expanded their climatic niches along PC2. Tree-shrub forms have expanded their climatic niche along all axes. Evolution of the climatic niche of ecomorphs along PC1 (b), PC2 (c) and PC3 (d). ANC of the ecomorphs are depicted on nodes. A dominant pattern of climate niche evolution is not evident along PC1 but a slight correspondence is shown in PC2 and PC3. Rock-boulder forms show a pattern of evolving towards extreme temperature conditions while a single species of Tree-shrub ecomorph, *Pseudophilautus regius* seems evolving towards extreme warm and dry conditions as well (see PC2 and PC3).

## Notes

### Competing Interest Statement

The authors have declared no competing interest.

